# Reconciling the tempo and timing of eukaryote emergence with clade dynamics and the fossil record

**DOI:** 10.64898/2026.03.25.714154

**Authors:** Corentin C. Flament-Loron, Niall Rodgers

## Abstract

The timing of the last eukaryotic common ancestor (LECA) remains a fundamental question in evolutionary biology and palaeontology. Fossils with morphologies compatible with a eukaryotic-grade affinity are reported as early as 1780 Ma, but no crown-group supergroups are confidently identified before the end of the Mesoproterozoic (ca. 1050 Ma). Recently, the late LECA hypothesis suggests that this absence of crown-assignable fossils implies a late Mesoproterozoic origin of the crown. Here we evaluate the compatibility of various LECA time frames with the evolutionary dynamics of the eukaryotic clade, the limited constraints of the fossil record and modern diversity. Studying stem-crown dynamics based on a birth-death model, we show that there is a tension between a long stem and the diversification rates implied by the size and age of the eukaryotic clade. We analyse the parsimony, implications and requirements of Late and Early LECA in terms of both modelling and the fossil record. We find that early LECA is parsimoniously consistent with diversification dynamics while a late LECA scenario requiring a long stem is less likely (but not impossible) under the utilised diversification dynamics. Finally, this study also highlights the importance of considering clade dynamics together with fossil evidence and molecular clock estimates when investigating the early evolution of life.

## Introduction

Eukaryogenesis marks the fundamental division of life^1^ that later led to complex multicellular organisms^2^ and modern ecosystems. The absence of a clear timeline for eukaryogenesis leaves a critical gap in connecting biological innovations to Earth’s environmental history^3,4^. The molecular clock estimates for the age of LECA have produced estimates ranging from 1000 Ma to 1900 Ma^5–9^, reflecting uncertainty in substitution rates and calibration points. Meanwhile, the fossil record may provide minimum ages for the emergence of stem and crown eukaryotes, but suffers from preservation, sampling and interpretation biases^10^.

Recent deep-time microfossil discoveries from rocks as old as ca. 1780 Ma present morphological characters compatible with a eukaryotic-grade cellular complexity that may indirectly evidence LECA characteristics, such as a cytoskeleton and an endomembrane system^11–15^. Their depositional distribution can also indirectly suggest the presence of mitochondria^16^. However, organisms closely resembling LECA, both morphologically and functionally, can still fall outside the crown group, for example, on a late diverging stem branch or by belonging to an unrelated complex prokaryotic lineage^3^. Indeed, extant representatives of such lineages (e.g., Asgard archaea) are now known to possess cytoskeletons and endomembrane systems and were possibly performing aerobic respiration^17–19^. Nevertheless, the preservation potential of these prokaryotes needs to be demonstrated, and evidence of recalcitrance for these morphological characters has been, so far, restricted to eukaryotes^20^.

Instead, a confident assignment to the crown group requires the identification of apomorphies associated with post-LECA eukaryotic supergroups^3,14^. No unambiguous crown super-group members have yet been reported before the end of the Mesoproterozoic^21–26^, despite earlier candidates^13^.

Based on these long-lasting uncertainties in fossil interpretations, the absence of crown eukaryote biomarkers, loose molecular estimates, and the young age of reported crown members, authors suggested an alternative scenario for the origin of LECA^3,6,8^: a late emergence with the crown eukaryote origin at the end of the Mesoproterozoic (1000–1200 Ma). This scenario implies that the late Palaeoproterozoic and most Mesoproterozoic eukaryotic-grade record consists of stem eukaryotes only (“long stem” scenario), or even of organisms not part of the total-group at all.

Practically, the late LECA hypothesis could be rejected if an early LECA hypothesis is validated, i.e., if unambiguous crown-eukaryote fossils or signatures are found in rocks much older than the late Mesoproterozoic. However, what has not been tested is the theoretical compatibility of each of these hypotheses and scenarios with the evolutionary dynamics of the eukaryotic clade itself.

Work in stem-crown dynamics has shown that the relationship between total group age, diversification rate, and crown emergence can be analytically approximated using birth-death models^27,28^. Birth-death processes model the stochastic dynamics of speciation and extinction and are used to answer many questions in macroevolution (e.g.,^29–31^).

To understand the timing of eukaryote evolution, any viable evolutionary model must account for the supergroup members of the late Mesoproterozoic as the only available direct minimum age for the crown group for now. The domain cannot be younger than this minimum age. On the other hand, the constraints posed by organisms bearing eukaryotic-grade traits in the late Palaeoproterozoic are much softer considering the challenge of placing them within the crown, the stem or even within the total-group^3^. Especially, considering the dual nature of the eukaryotic stem before the endosymbiotic merger^4,9^. Hence, these fossils can only provide a soft minimum age for the total group. Finally, the model must be consistent with the living diversity of eukaryotes, and ideally consistent with other frameworks, like molecular clock estimates.

Here, we use a birth-death framework to assess the compatibility of stem-length, the fossil record, diversification rates and living diversity. We specialise the framework of Budd and Mann ^28^, which used birth-death modelling to study the generic dynamics of the emergence of crown groups to the study of the emergence of eukaryotes, incorporating the additional effects of stochasticity that should be considered^32^. We note that the size and timescales of the eukaryotic emergence make them an outlier case in clade dynamics. Our goal is not to precisely model the diversity of eukaryotes over time, nor are we trying to provide a precise age for crown or stem emergence. Simply, we wish to test the required evolutionary dynamics of LECA emergence scenarios, using the simplest, most parsimonious model, and analyse the trade-offs and assumptions inherent in each scenario.

Our results highlight tension in scenarios involving a long stem. Long stems are highly improbable (but not impossible) under the constraints of very large and ancient clades whereas short stems are more favourable under the evolutionary dynamics of large and ancient clades. However, both scenarios imply questions about the current and future constraints provided by the fossil record for which we account herein.

If crown eukaryotes arguably emerged early, possibly consistent with (but not dated by) the Palaeoproterozoic record of complex microfossils, this record may contain crown members that, if preserved, have not yet been unambiguously identified.

## Short review of empirical constraints

### Minimum age for the total-group, *T*

The origin of total-group eukaryotes began with the emergence of at least two first eukaryotic common ancestors (FECAs), nuclear FECA and mitochondrial FECA^4^, which diverged from their respective archaeal and bacterial sister groups at different times^9^. This means that the total-group eukaryote is not represented by a single diverging node. It comprises two independent stem lineages that merged in mitochondrial endosymbiosis, after which a single lineage continued to the last eukaryotic common ancestor^4,9^. Because only one descendant lineage of this merger survives, the event does not correspond to a node in phylogenies, and therefore cannot be directly dated by molecular clocks^4^.

Some early microfossils present morphological or ultrastructural characters that indirectly suggest the presence of a complex cytoskeleton, endomembrane system, and complex enzymatic toolbox^3,8,14,15,20,33^. However, these characters are insufficient to clarify stem or crown position within eukaryotes^3,14^ and similar traits are found in sister lineages of complex prokary-otes^17–19^ (although possibly not recalcitrant enough to be preserved^20^). Taken together, the evidence shows that aspects of the cellular complexity relevant to eukaryogenesis were already present by the late Palaeoproterozoic.

Furthermore, the archaeal host was already cytoskeletally complex before endosymbiosis^9^, meaning fossils bearing these traits could sit on the host stem before or after the merger. Relevant fossils include specimens from the 1780 to 1129±4 Ma McDermott Formation, McArthur Basin, Australia^15^, from the 1744±22 to 1411±27 Ma Ruyang Group, Shanxi Province, North China Craton^11,34^, from the 1673±10 to 1634±9 Ma Changzhougou and Chuanlinggou formations^12,13^, and from the *>*1642±3.9 Ma late Palaeoproterozoic Limbunya Group, Birrindudu Basin, Australia^14^.

Given the above, the age 1780 Ma represents a soft minimum age for the total-group, conditional on these fossils documenting innovations on the lineage leading to LECA. If they fall outside the total-group, the constraint applies instead to a deeper node. In either case it bounds the total-group as a whole, not either FECA individually.

It could potentially be argued that these fossils also provide a maximum constraint for LECA emergence, because the cellular characters of LECA are expected to have appeared by the time of crown emergence. However, this is directly conditional on the current fossil record, and only valid until the discovery of new older complex fossils.

Other older possible, but more ambiguous^3,15^, candidates include the coiled macrofossil *Grypania spiralis* from the 1891±3 Ma Negaunee Iron Formation, USA^35^ and large spheroidal microfossils reported from the 3.2 Ga Moodies Group suggested to be possible stem eukaryotes^36,37^. Aside from their size, these organisms lack traits that could conspicuously be linked to eukaryotic-grade cellular organisation.

In the mathematical framework introduction, we discuss the notion of total-group and the additional nuances that might apply in the context of reconstructing eukaryote clade dynamics with birth-death models.

### Minimum age for the crown-group, *t*_**cg**_

In contrast to the total-group, minimum age for the crown-group can be more directly inferred from fossil evidence. Unambiguous crown group members are found from the late Mesoproterozoic and include representatives of various total eukaryotic supergroups, possibly crown, based on specific, clade-related morphological, ultrastructural and molecular traits. They include *Bangiomorpha pubescens* (1047 +13/–17 Ma), possibly a member of the Bangiales (Rhodophyta, Archaeplastida)^21,22^; *Protorocladus antiquus* (ca. 950-1050 Ma), a total Chlorophyta (Archae-plastida)^23^; *Arctacellularia tetragonala* (ca. 950-1030 Ma), a total Archaeoplastida^24^; *Ouras-phaira giraldae* (1013 ± 25-892 ± 13 Ma), a possible total Fungi (Opisthokonta)^25^; *Ouroboros soroboruo, O. vulgaris* and *O. torus* (1078 ±24 Ma), possible euglinids (Discoba)^38^; and 994 ± 48 Ma *Bicellum brasieri*, a suggested Holozoa (Opisthokonta)^26^. An older candidate is *Qingshania magnifica* from the 1634±9 Ma Chuanlinggou Formation, a multicellular, possible archaeplastid^13^.

A conservative estimate for the minimum age for the crown group is therefore 1050 Ma. The late LECA hypothesis^8^ suggests that the crown might have emerged shortly before that minimum limit.

### Living eukaryote diversity

We consider the end product of our model to be the living diversity in species, representing the number of extant lineages at the present. Currently, approximately 2.5 million species of eukaryotes are catalogued in the Catalogue of Life^39^. However, in this cataloguing, microbial species are strongly under-represented. In any case, this is only the observed diversity and the actual living diversity may be much higher. Example estimates place it at 5 ± 3 million species^40^ or 8.7 ± 1.3 million^41^. Hence, we assume the conservative value of 2.5 million present lineages.

## Mathematical framework

### Mathematical modelling of Birth-Death processes

There is a long history of using mathematical modelling to understand evolutionary dynamics^27–30,42^. In this work, we aim to use standard, and intentionally conservative, results of birth-death modelling^28^ constrained by the modern eukaryotic diversity and interpretation of the fossil record to examine scenarios for the timing of the emergence of LECA.

In this work, we assume constant speciation and extinction rates, as is standard for the field^28,30^. Time-varying rates enter into unconditioned quantities such as the expected number of species via an integral of the net diversification rate over time^30^, and for such quantities this is equivalent to a constant rate equal to the time average. Conditioned quantities do depend on the individual time-varying rates; however, we discuss rate changes and mass extinctions, in the feasibility section and argue that these caveats require additional assumptions which are not constrained and may have marginal impact on the broad conclusions. Another issue with changing to a more complex model is the lack of constraints on the variables and form of the time variation. Given this, we explore the behaviour in a parsimonious model with which we can sweep the whole parameter space. It is also important to note the difference between something being improbable for a typical case under a stochastic model and unlikely under any case in the stochastic model^32^, which we explore below.

In this work, we adopt the standard notation of the field following Budd and Mann ^28^ . We take the per-lineage speciation rate *λ* in My^*−*1^ and a per-lineage extinction rate *µ* in My^*−*1^. These can be combined to give a diversification rate *r* = *λ* − *µ* and a turnover rate 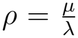. In this work, we focus on growing clades so we enforce *r >* 0 and *ρ <* 1. We take the convention that time runs forward to the present *τ* ∈ [0, *T* ]. With the key dates running backward from the present, the total group age, *T* and the crown age, *t*_cg_. We also define the length of the stem as *g* = *T* − *t*_cg_.

It is important to note that birth–death models describe a bifurcating process of lineage diversification. In Budd and Mann ^28^, the stem process is considered from an initial point of origin to the first bifurcation ancestral to lineages that survive to the present, corresponding to the total-group and crown-group origins, respectively. However, in the case of eukaryotes, the point of origin in the model, *T*, does not necessarily correspond to the actual evolutionary origin of the total group. Rather, it would represent the point at which the lineage begins to evolve according to the bifurcating process described by the model. In the case of eukaryotes, it may be more biologically appropriate to associate this point with the establishment of a single, independently evolving lineage following the endosymbiotic merger, rather than with the earlier divergence of its prokaryote ancestors and the reticulated evolution during the endosymbiotic event(s) (see Discussion). Nevertheless, for simplicity and consistency with the terminology of^28^, we retain the designation “total-group *T*” throughout the derivations and modelling below.

An important simplifying limit is *rT* ≫ 1, appropriate for large and ancient clades (clearly eukaryotes would fall into this category). However, for generality we do not always work in this limit so we can evaluate all LECA scenarios. When *rT* appears it is usually as an exponent, therefore *rT* does not need to be very large for the approximation to be reasonably accurate. One may estimate that *rT* is roughly of the order ln *N* for a growing birth-death process, where *N* is the number of species at time *T*, meaning that *e*^*−rT*^ would be expected to be very small for a clade as large as the eukaryotes. As this work involves matching of scenarios some quantities may be conditioned on survival and some on crown formation however the numerical impact of the difference in these conditions is small for 2.5 million species and does not change our qualitative orders of magnitude conclusions.

### Probability of survival to the present

Following Budd and Mann ^28^, the number of species, *N*, that survive to the present for a homogeneous birth-death process without mass extinctions which begins *T* years in the past conditioned on the survival of the process is given by the conditioned geometric distribution which can be written as

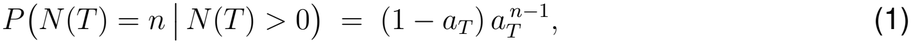

where

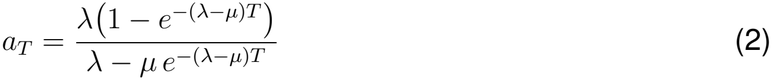

or equivalently

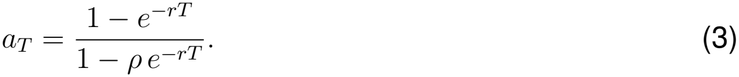

From Budd and Mann ^28^, the probability of a birth-death process beginning with one species at *T* surviving to the present can be written as

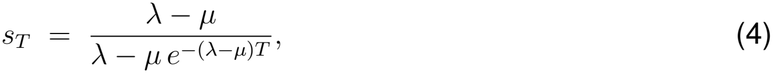

or equivalently

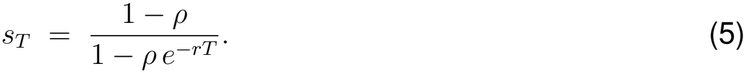

In the limit *rT* ≫ 1, this can be simply written as

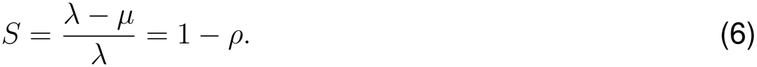

It can be shown that the expected mean number of species conditioned on survival of the process can be written

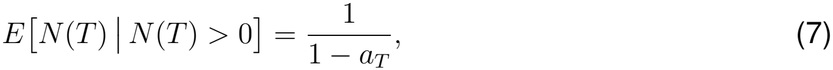

by using either the relationship between the conditioned and unconditioned means or via direct computation via geometric series and using the reordering of sums and derivatives. This can be equivalently written as

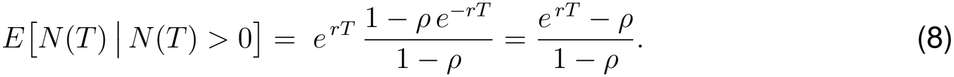

In the limit *rT* ≫ 1, this leads to the simple form (*S* also in the limiting form)

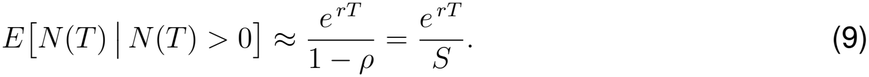

Additionally, we can write the probability of the survival of at least *k* species^28^

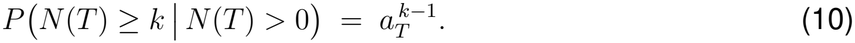

This probability is clearly very sensitive for large *k* and rapidly becomes very close to zero unless *a*_*T*_ is close to 1.

### Stem-Crown gaps with diversification rates

The cumulative probability of crown group formation within time *g* can be written, equation 11, using a result from Budd and Mann ^28^, conditioned on crown formation (as opposed to only survival) by elapsed time *T* . The cumulative distribution function (cdf) of this truncated exponential distribution gives the probability that the crown has formed within *g* elapsed time from the emergence of the stem and can be written as

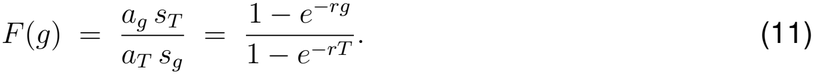

Intuitively, at *g* = 0, we have *F* (0) = 0 and at *g* = *T* we have *F* (*T* ) = 1. Additionally, this is independent of turnover rate. Using equation 11, we can find the probability density function *f* (*g*) by using

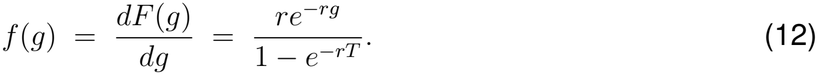

This allows us to find the mean and median values of the gap distribution by solving the following,

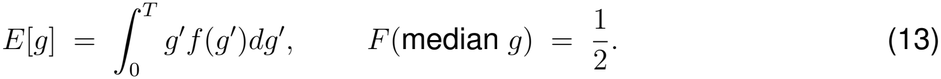

Hence, the mean and median of the gap distribution (equivalently described as time to crown formation) described by cdf 11 are

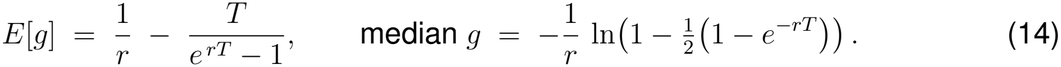

In the large ancient clade limit *rT* ≫ 1, the truncation is negligible and the average gap statistics are approximately exponential with rate *r*, so that

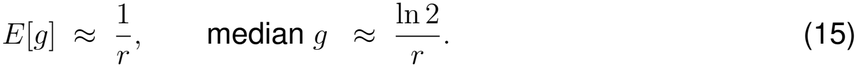

Taking the expression for the mean gap and writing in terms of the age of the total group and the age of the crown leads to an expression containing the diversification rate implied by the mean stem crown gap which depends only on the age constraints,

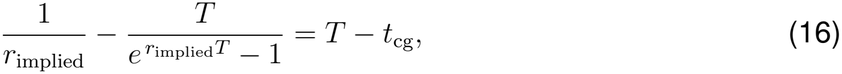

which can be written as

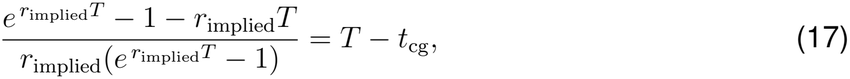

which demonstrates that the gap is always positive for *r*_implied_ *>* 0. This cannot be solved analytically for *r*_implied_ so it must be solved numerically. However, it can be shown analytically, see appendix B, that the left-hand side of equation 16 is strictly decreasing for positive *r*_implied_ with unique positive solutions existing for 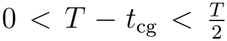. In the large and ancient limit *rT* ≫ 1, equation 16 reduces to *r*_implied_ ≈ 1*/*(*T* − *t*_cg_). Note that in all figures we use the full numerical solution as this does not hold in all late LECA regimes and would give unfeasible answers for small *r*_implied_.

Of course, the mean implied values do not fully capture the dynamics^32^ of the origin of a crown and it is important to consider the tails of the distribution and assess the likelihood of a given gap formation when considering the full distribution. *r*_implied_ tells us that given a proposed stem length what would an average diversification rate look like, not the full possibility space of diversification rates. Using equation 11 to write the probability of a gap greater than *g* leads to

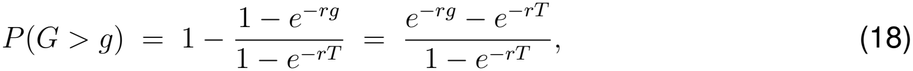

which in the limit of *rT* ≫ 1 when *g* is not close to *T*, is approximated by

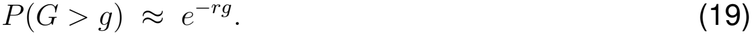

From equation 19, we see that the probability of large gaps decreases exponentially in *rg* hence large gaps are disfavoured under fast diversification. However, exponential distributions can have long tails (depending on the parameters) so lower probability long gaps cannot be fully ruled out. In fact, more generally the fact that birth-death models result in exponential distributions which do not have sharp decay fundamentally limits the precision of statements that can be made when analysing historical processes. We discuss the methods to solve equations numerically and plot the results in Appendix A.

## Results

### Mean stem lengths and implied diversification rates

Firstly, we analyse the typical diversification rates implied by a mean gap. This is plotted in figure 1 which shows the numerical solution of equation 16.

**Figure 1:**
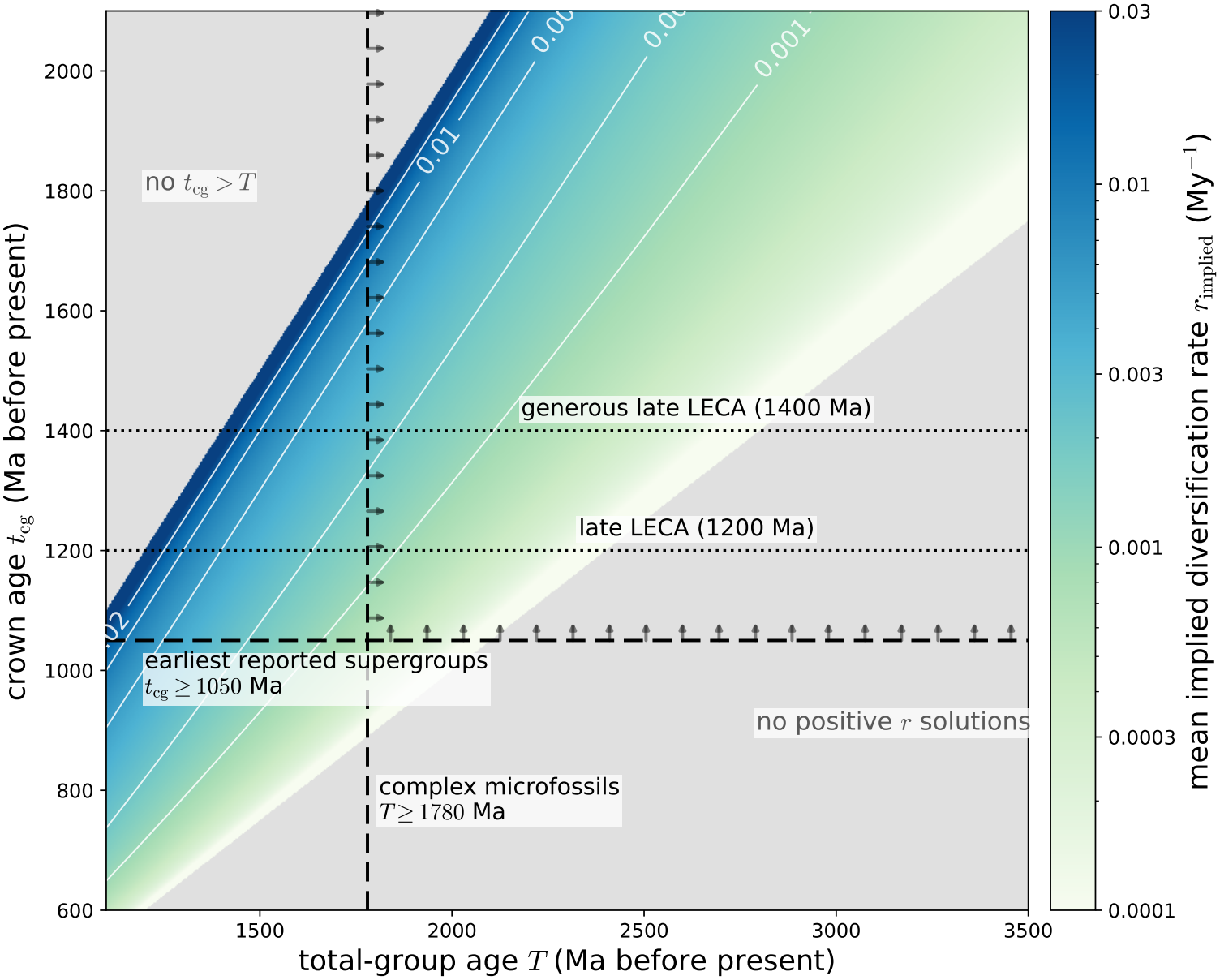
Implied diversification rate across LECA scenarios plotted via numerical solution of mean-gap equation 16 with allowed solutions shown as a heatmap. Overlaid fossil bounds are described in the main text with potential LECA scenarios overlaid for reference. Arrows point towards fossil-feasible region.

We see that figure 1 has excluded regions. This is due to the fact that we cannot have a crown older than the total group and that mean-gaps greater than 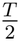 require a negative rate which is not feasible. We can see that in this mean-implied framing long gaps such as LECA emergence just before the crown at 1050 Ma and assuming the total bound at 1780 Ma suggest a rate well below 0.001 My^*−*1^. When the time of emergence of LECA is moved back, keeping the same total group age, rates become more feasible. In general, we see that reasonable rates are suggested by short gaps and longer gaps imply very slow rates which are hard to reconcile with the extinction risk of having very low rates for hundreds of millions of years.

### Scenario correspondence with modern diversity

Rather than simply analysing rates implied by a given mean gap, we can look at how these implied rates interact with modern diversity or how the modern diversity can be used to suggest rates in the reverse direction. To find what mean rate is suggested by modern diversity we can simply solve equation 8 and equate it to modern diversity, *N* = 2.5 × 10^6^,

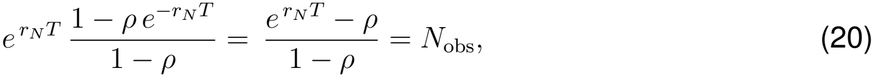

which has solution

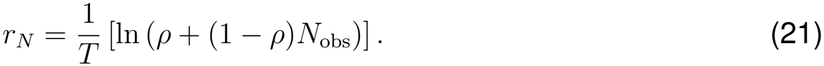

For very large clades (with *ρ <* 1), we have *r*_*N*_ ≈ ln *N*_obs_ + ln(1 − *ρ*) */T* as the contribution of turnover is small relative to the modern diversity. This means that roughly *rT* ∼ ln *N* so the size of modern diversity can give an indication that large *rT* -regime applies or not. Equation 21 is not a full maximum likelihood result conditioned on exactly *N* species, but a moment matching exercise to compare “typical” scenarios.

#### Probability of observing given gaps

Equation 18 gives the probability of observing a gap greater than a given value as a function of the age of the total group and the diversification rate. Using equation 21 as the rate implied by modern diversity, one can then write the probability of observing a given stem-crown gap greater than *g* by substituting into equation 18. This leads to

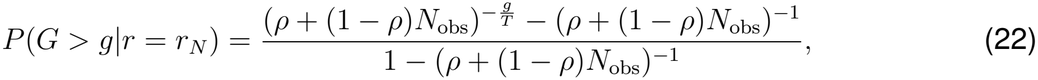

which simplifies for large *N* (and fixed 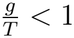) to

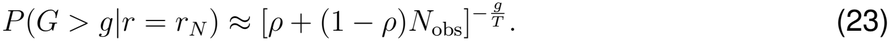

Equation 23 gives a good sense of what controls the probability of given stem lengths. The length of the gap enters as a ratio of the total group age which makes sense when we have eliminated *r* in this way. This is clearly intuitive as the probability of gap of a given length is really a question of what fraction of the time of the total group is spent before the emergence of the crown. The quantity again enters in the exponent, so we see fast decay but also long tails which are hard to rule out. Again intuitively, the probability of a long gap is suppressed as modern *N* grows, which requires fast diversification. The contribution of turnover is important but when discussing clades with millions of modern species it does not affect the qualitative trends that we observe. Equation 22 gives the probability of observing a gap greater than *g* using the rate implied by the modern diversity which is distinct from the probability of observing a gap greater than *g*^*′*^ conditioned on the modern diversity *N* .

In figure 2, we plot equation 22 for three representative turnovers across varied crown and total group ages.

**Figure 2:**
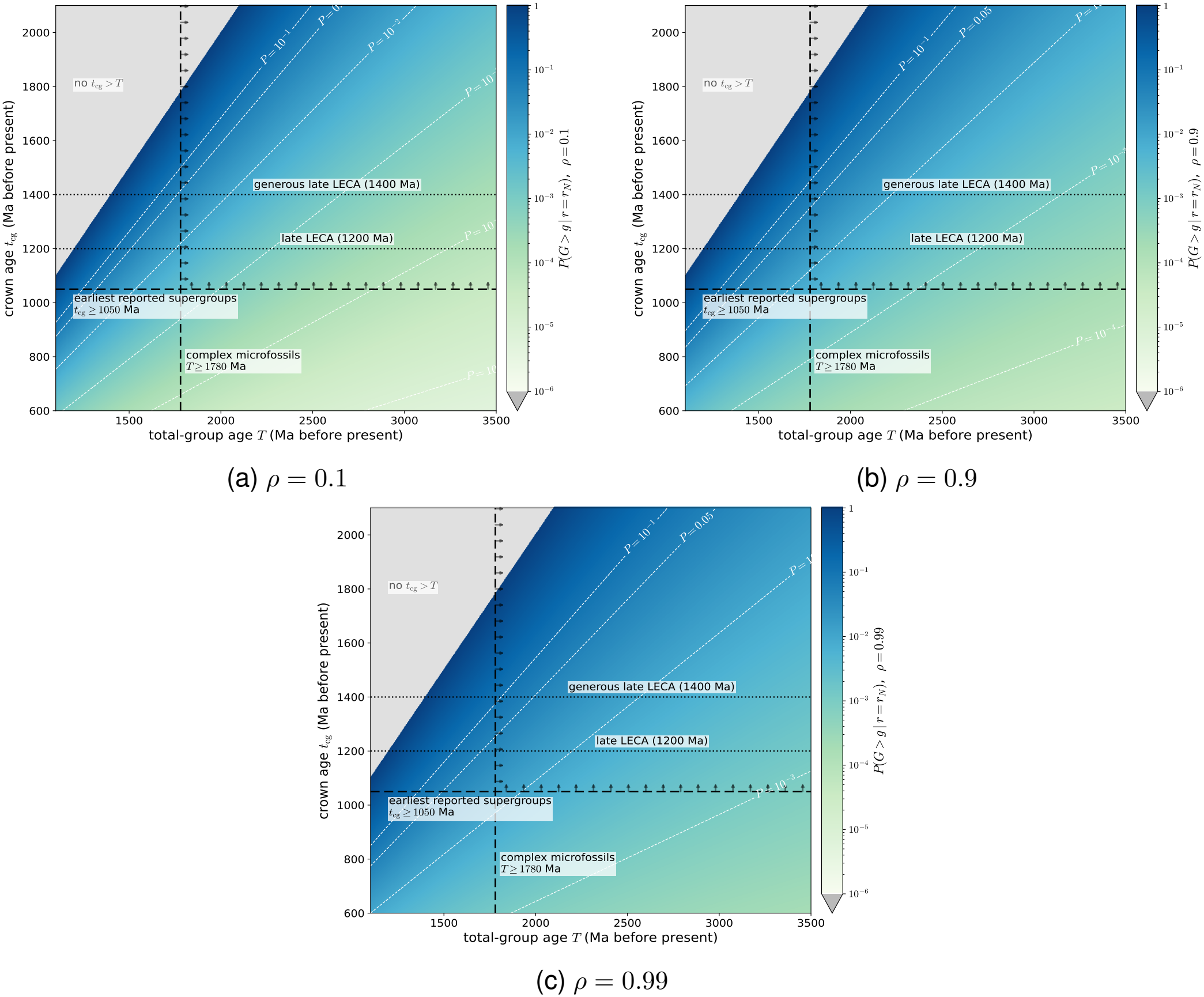
*P*(*G* > *g*|*r* = *r*_*N*_) across crown and total group ages. Each panel (a), (b), (c) gives a representative turnover *ρ* = 0.1, 0.9, and 0.99. Overlaid fossil bounds are described in the main text with potential LECA scenarios overlaid for reference. Probability values under 10^−6^ are set to zero for legibility. Arrows point towards fossil-feasible region.

In figure 2, we observe that probabilities are near one when *T* and *t*_cg_ are similar as nearly all gaps will be above zero. As the stem gets longer the probabilities decrease but due to the long tail there is still a very small chance to observe gaps of this magnitude. The contours give the probability of observing a gap greater than a given value under the model and we see that short stem scenarios are by far the most probable.

#### Modern diversity using mean-gap implied rates

It is also possible to get a qualitative sense of the likelihood of various LECA scenarios by taking the rate implied by the mean gap and then using this as an input into calculations which predict or depend on modern diversity. This is a stronger condition as we are essentially asking if an outcome is compatible with typical inputs rather than whether this is possible in general given the full probability distribution. We can input the mean gap implied rate, *r*_implied_, into equation 8 to give the expected modern diversity of species and into equation 10 to give the probability of observing at least modern eukaryotic diversity under this rate. This is shown in figure 3. As this is not independent of turnover, we plot three different turnovers; however, despite the impact of turnover on the absolute values being strong, it does not affect the qualitative conclusions. Figure 3 should be read as separate plug-in calculations rather than a joint probability calculation conditioned on diversity and observed gap.

**Figure 3:**
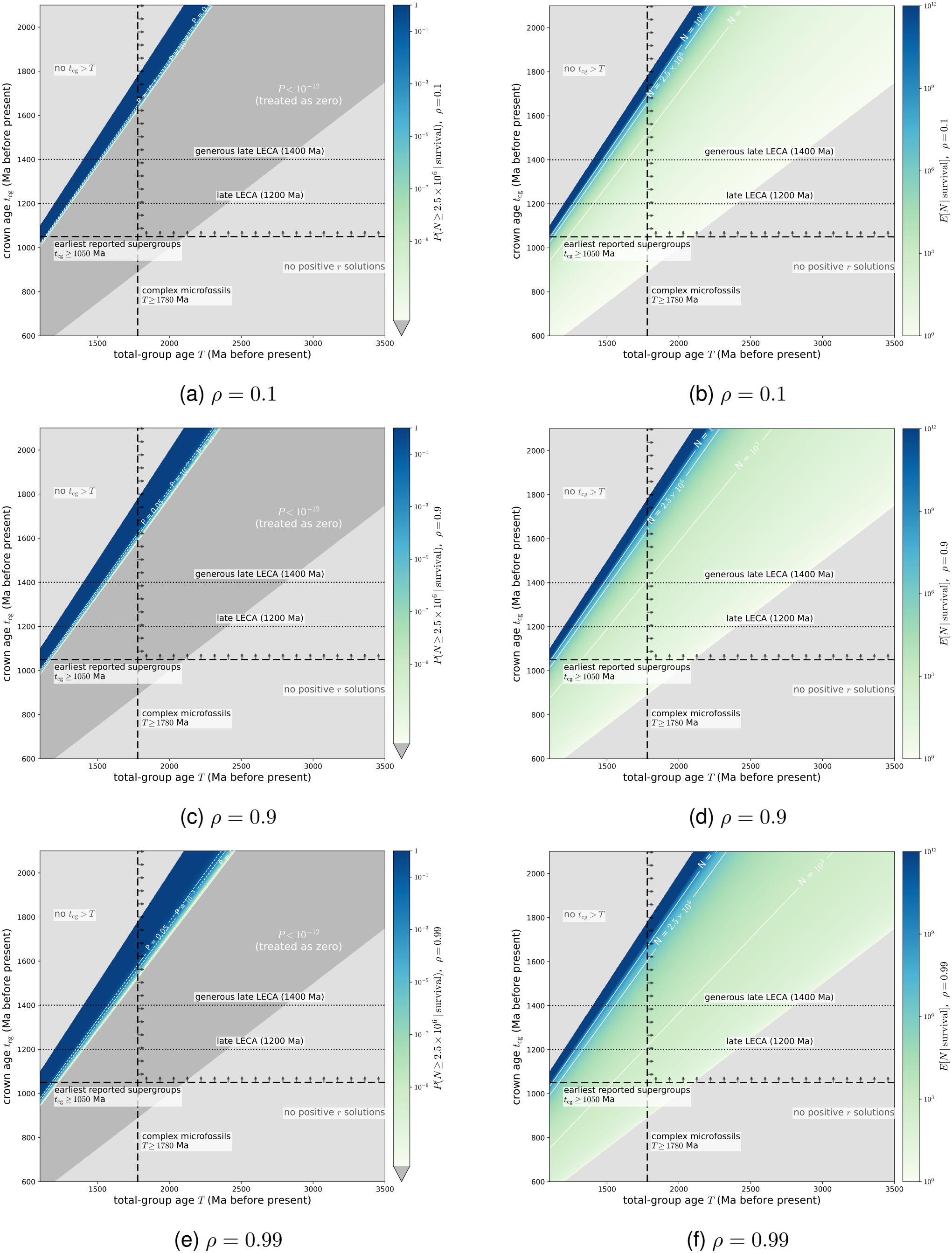
Diversity tests at the gap-implied mean rate using various LECA scenarios with varying turnover *ρ* = 0.1, 0.9, and 0.99. Probabilities of at least modern diversity under the mean implied rates are given in the left column [panels (a), (c), (e)] while expected species number is given in the right column [panels (b), (d), (f)]. Overlaid fossil bounds are described in the main text with potential LECA scenarios overlaid for reference. Probability values under 10^−12^ are set to zero for legibility. Arrows point towards fossil-feasible region.

The left column of figure 3 shows that under mean-gap-implied-rates with any long stem the probability of observing modern diversity is vanishingly small. Due to the large size of the eukaryotic clades at this mean value it is very unlikely to observe a modern clade of the order of millions of species using this comparative mode. A similar result is shown in the right column of figure 3 where the expected number of species is only a handful or a few hundred for all long stem scenarios, orders of magnitude below the modern diversity.

When *rT* ≫ 1 the expected gaps are given by equation 15. The expression for the median gap, equation 15, is equivalent to the approximation which relates the diversification rates and stem lengths found in Budd and Mann ^27^, their equation 18. Using this (or deriving from equation 19), we can write

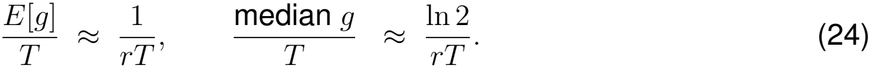

Then using that for *r*_*N*_ *T* = ln (*ρ* + (1 − *ρ*)*N*_obs_) as the rate implied by modern diversity we find that we can write

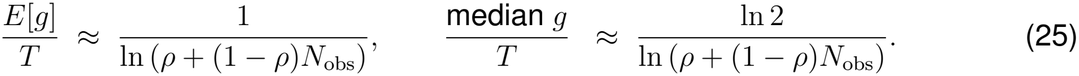

This means that again in general, it is expected that for very large, diverse clades that the length of the stem is a small fraction of the age of the total group. Again, subject to the limits of describing an exponential distribution via its mean^32^.

### Implications of the mathematical framework for LECA constraints

The core implications of the mathematical framework are as follows. If one makes parsimonious constant mean-field rate assumptions and does not allow for external factors, one generally expects that a “typical” ancient clade which has a large modern diversity should have a short stem because of the tension of a long stem (requiring low diversification rate) with high modern diversity (requiring higher diversification rate). Similar conclusions have been reached by others working on birth-death modelling in more generic contexts such as^28,43^. However, one must also note that birth-death models depend on exponential distributions which may have a long tail^32^, so long stems might happen to be a low probability (exponentially suppressed) draw from the distribution. This framework suggests the following before considering fossil constraints and external factors: one may have an early total group and early LECA, a late total group and a late LECA, but an early total group and late LECA is unlikely (although not impossible) in the absence of external factors due to the discrepancies in the required rates.

## Discussion

### Feasibility, possible scenarios, and external factors required

In this section, we describe possible external factors and modelling omissions along with their directionality and potential impact before commenting on the feasibility of various scenarios in light of their requirements and support from the fossil record.

#### Influence of possible additional factors

An obvious factor affecting the gap between the origin of the crown and the total group is the possibility of mass extinctions^28^. Budd and Mann ^28^ state “Typically, the stem group diversifies rapidly until the crown group emerges, at which point its diversity collapses, followed shortly by its extinction. Mass extinctions can disturb this pattern and create long stem groups such as the dinosaurs. Crown groups are unlikely to emerge either cryptically or just before mass extinctions, …”.

Mass extinctions can extend the length of the stem by eliminating early-diverging lineages and delaying the establishment of the crown-group; however, they do so at the cost of suppressing diversity, so they do not obviously resolve the discrepancy of a long stem in a highly successful clade. For a mass extinction to affect the stem length it must also be large^28^. At the end of the Archean and early Palaeoproterozoic, major environmental perturbations associated with the Great Oxidation Event and Palaeoproterozoic glaciations may have caused losses of microbial diversity. However, the concept of mass extinction is difficult to apply to microbial ecosystems, where species boundaries are poorly defined and where profound ecological turnover does not necessarily entail the extinction of phylogenetic lineages^44,45^. We cannot exclude the possibility of substantial microbial extinctions. However, it is also unclear whether, if such events did occur, they would be sufficiently severe, have the necessary phylogenetic consequences, and occur at the appropriate time to prolong eukaryotic stem(s).

Even if we allow for a large stem-extending mass extinction, this effectively resets the clade diversity at the extinction point, requiring higher rates after the event in order to reach modern diversity. This scenario presents an unfavourable sequence of requirements: first that high rates are possible after eukaryogenesis but that they initially do not occur as we have no fast crown emergence; then a mass extinction occurs; and then fast rates do kick in and we reach modern diversity. We do not have evidence for each of these requirements and the mass extinction ends up playing a secondary role to the rate change in this scenario.

One possibility to make an early total-group origin and late LECA work would be for the diversification to initially be very slow and then speed up at the time of LECA. This contradicts what would be expected of a large clade^28,43^ and it is unclear what effect, if any, one expects endosymbiosis and other evolutionary changes should have on rates. One may hypothesise that some combinations of specific environmental factors suppressed the diversification of early eukaryotes on the early Earth such as limited oxygen availability, lower rate of primary productivity, nutrient scarcity, or restricted ecological opportunities^8,10,46–48^.

However, the required change in rates would be substantial. A hypothetical long stem with total group at 1800 Ma and LECA at 1100 Ma would imply (based on numerically solving for the exact mean-implied gap) an approximate rate of 0.0008 My^*−*1^. In contrast, starting from a single species (starting at two species gives a similar result) at the origin of the crown group to arrive at expected modern diversity (setting turnover to zero) implies a rate of 0.0134 My^*−*1^ which would be a roughly 17-fold increase. This rate (approximated from two separate models) of increase, although not impossible, would need an additional justification for its occurrence. All these calculations should be caveated as rough estimates depending heavily on modelling assumptions; small changes to dates, number of starting species and turnover affect the values, which is why we adopt the conservative substantial but not impossible conclusion.

Of course, as pointed out by^32^ it is possible to observe long stems even in conditions in which you may not expect them on average simply due to the stochastic nature of the model. In the particular case of eukaryotes the size and age of the clade make long stems less likely. However, fundamentally we have a sample size of one for the origin of eukaryotes so one cannot rule out that perhaps the eukaryotic origin on Earth was atypical and far from the mean.

One possible area of model error would be errors in the present diversity^39^. This is not as serious a concern as it may appear due to the fact that roughly one expects *rT* ∼ ln *N* meaning that the impact of errors in diversity on required *rT* is suppressed by the logarithm. Underestimation of modern diversity is, purposely on our part, likely^40,41^ and larger modern diversity would impose higher rate requirements, making a longer stem less likely.

The push of the past and the pull of the present^27^ are well known survivorship biases that can play a role in birth-death modelling. Pull of the present affects the reconstructed diversity curves as reconstructions approach the present due to extinctions not yet occurred^27^. This is not a concern for our question, as we directly sample living species. We likely underestimate the total number of living eukaryotes from a practical sampling perspective and errors in modern diversity make a small contribution, as discussed above. The push of the past comes from conditioning on survival and results in faster observed speciation rates near the origin of a clade^27^, which is already part of our model as we work with conditioned quantities on long timescales. This would not be an additional effect to change the conclusions with additional conditioning on large modern diversity further suppressing long stems.

#### Early total group and early LECA

This scenario is consistent with diversification dynamics and is likely under standard birth-death models, as a short interval between total-group and crown-group origin is compatible with a high extant diversity. Palaeontologically, this scenario is also consistent with the fossil record of organisms at the end of the Palaeoproterozoic presenting cellular and ecological characters compatible with a eukaryotic cellular complexity^11–15,34^. In this scenario, the substantial interval between LECA and the earliest unambiguously identified crown-eukaryote fossils in the late Mesoproterozoic remains to be explained, but this interval likely reflects preservation, sampling or detection and interpretation biases of the existing Mesoproterozoic record^8,10,49^. The potential future confirmation of early crown candidates, such as *Qingshania* as an archaeplastid in the late Palaeoproterozoic^13^, will considerably strengthen this scenario.

#### Late total group and late LECA

This scenario is in principle compatible with birth-death model expectations (relatively short total-crown interval). However, a late origin of the crown-group leaves less time for the accumulation of modern eukaryotic diversity and therefore requires higher diversification rates. This scenario is also potentially inconsistent with the late Palaeoproterozoic complex microfossils, even with their phylogenetic affinities remaining uncertain. Moreover, this scenario is difficult to reconcile with molecular clock estimates supporting older divergence times^9^.

#### Early total group and late LECA

This scenario is less likely under the expectation of evolutionary dynamics^28^, as it implies a prolonged total-crown interval in a highly diverse clade. However, a low-probability stochastic diversification cannot be ruled out^32^.

The work of Budd and Mann ^27^ argues that stem groups are usually characterised by high observed diversification rates and low diversity whereas crown groups show low diversification rates and increasing diversity. Therefore, the pattern observed in the Precambrian fossil record does not necessarily reflect the true timing of diversification. Indeed, the Mesoproterozoic record contains steady reports of fossils with a eukaryotic-grade cellular complexity^10,49^, but these reports are highly correlated with the number of sampled assemblages, introducing a consequential bias in diversity reconstructions^10^.

Instead, to accommodate this scenario (without a low-probability event), a prolonged period of low net diversification, an extensive extinction of stem lineages, and large changes in rates could be considered but require many additional assumptions.

In addition, this scenario is compatible with the interpretation of late Palaeoproterozoic fossils as stem eukaryotes but will be challenged if the early supergroup crown-candidates were to be confirmed as crown (e.g., *Qingshania*).

In molecular clock reconstruction, a scenario with an early nuclear FECA and a much later LECA (ca. 1.3 Ga gap in^9^) is presented. However, this interval may partly reflect evolutionary processes preceding or accompanying mitochondrial acquisition, rather than branching diversification within a single eukaryotic lineage from the start. Moreover, this estimate for LECA is still considerably older than a late Mesoproterozoic origin.

### Further discussion

We intentionally select a simple model given the low data regime in which we are working. A more complex alternative may be to incorporate diversity-dependent diversification (DDD)^50,51^, however it was argued in Budd and Mann ^28^ that these models are more likely to show rapid crown expansion than the homogeneous case. We acknowledge the limitations of describing a long evolutionary history with a simple homogeneous birth-death model^43^, in particular to describe the behaviour of a successful outlier clade that may not behave as the average clade. However, note that the key argument of this work is the statistical tension between a long stem and the fast initial diversification implied by the existence of a modern large clade rather than precise predictions. More complex models such as the Covariant Evolutionary Tempo model^43^ also support this fast initial diversification with Budd and Mann ^43^ stating “*these large clades are expected to be characterised by explosive early radiations* …”.

The reticulate nature of eukaryogenesis raises important questions about the biological interpretation of the point of origin in birth–death models applied to eukaryotes. The mathematical framework used here describes diversification from an initial lineage to the emergence of the first bifurcation ancestral to surviving lineages that we interpret here as the origin of the eukaryotic crown group. Because eukaryogenesis involved the integration of archaeal and bacterial ancestors, the divergence of these ancestral lineages, notably nuclear FECA, does not necessarily correspond to the establishment of a single, independently diversifying eukaryotic lineage. Instead, the point of origin in our model could be interpreted as the establishment of such a lineage following endosymbiosis, when its evolutionary history became more appropriately approximated by a bifurcating process as opposed to a reticulate process before that^52^. Under this interpretation, the relatively short origin-to-crown intervals favoured by our framework could provide constraints on the timing of the establishment of the post-symbiotic eukaryotic lineage relative to LECA. Combined with independent molecular-clock estimates of LECA, these results could therefore help constrain the timing of this evolutionary transition, otherwise undatable with node-based phylogeny. Although this transition need not coincide precisely with mitochondrial acquisition, it may provide indirect information about the timing of the endosymbiosis, provided that the interval between mitochondrial acquisition and the establishment of an independently diversifying lineage was relatively short. Recent molecular clock estimates by^9^ suggest a LECA age of 1.80–1.67 Ga. Our simple model then roughly suggests a most probable endosymbiosis on the order of a few hundred million years before this time frame, with the caveat that endosymbiosis and the start time of the model may be offset and that longer intervals cannot be ruled out under the model.

We expect the methodology of this work and conclusion, that long stems are disfavoured in large modern clades^28^ (subject to stochastic fluctuations^32^) to hold quite generally, as this does not require eukaryotic specificity. However, the eukaryotes are an exceptional case, given their size and the long timescales involved.

These results also support some molecular clock estimates for the age of LECA. Estimates span more than one billion years, reflecting fundamental disagreements in calibration, substitution rate, and fossil assignments. Older age estimates, placing LECA at 1600–1900 Ma, rely on Proterozoic calibrations such as *Bangiomorpha pubescens*, previously dated at 1198 ± 24 Ma and now at 1.047+0.013/–0.017 Ga^22^. The inclusion of *Bangiomorpha* (with its previous age) seems to consistently pull the crown age estimates older^5,7,8^. Younger age estimates emerge when Proterozoic calibrations are excluded or when *Bangiomorpha* is reassigned to total-group red algae rather than crown-group red algae, a choice justified by the general body plan and anatomy of the fossil^6,8^. However, recent work dating the timing of eukaryotic gene duplication provides a new estimate of 1800–1670 Ma for LECA^9^.

Regardless of absolute age, molecular estimates^5,7,9^ indicate a rapid diversification of the eukaryotic supergroups after LECA, within 300 Ma. This is consistent with the record of organisms compatible with eukaryotic-grade cellular complexity in this time interval, where the record is demonstrably incomplete but actively studied with supergroup candidates being suggested^11,34,53–57^. An early rapid diversification in the late Palaeoproterozoic–early Mesoproterozoic is also compatible with recent estimates for the origin of Archaeplastida (the supergroup including red and green lineages), placing their crown origin at 1712–1387 Ma^58^, as well as with estimates for the origin of Amoebozoa, Excavata, Opisthokonta and SAR^5^ and suggestions of Proterozoic ecological diversification^59^. Rhodophyta, Chlorophyta, Opisthokonta groups, emerging later in eukaryotic phylogeny, would originate later than this interval, supporting their appearance in the fossil record at the end of the Mesoproterozoic^21–26^.

## Conclusion

In conclusion, we have analysed the compatibility of LECA origin scenarios with birth-death modelling and compared it with the main constraints provided by the fossil record. We find that the modern diversity of eukaryotes implies that a short stem, before the origin of crown eukaryotes, is the most likely scenario in the absence of additional evolutionary or ecological factors. Such a scenario is also compatible with known fossil constraints and recent molecular clock estimates for the age of the FECAs, LECA, and eukaryotic supergroups. However, we highlight that we only have one example in the history of life of the origin of eukaryotes and whether this is a high-probability or low-probability draw cannot be verified without more confirmation from the fossil record. Finally, our results use bifurcating birth-death modelling to analyse the eukaryotes, characterised by two stem origins and a resulting series of reticulated evolutionary events. In this evolutionary framework, such a model might be helpful to characterise the time of origin of the linear bifurcating process, which in eukaryotes corresponds to the aftermath of endosymbiosis.

## Data and code availability

This study is theoretical and all the data required for the calculations (the fossil ages and living diversity estimates) are provided within the main text. All results are a direct application of the equations mentioned in the text. The figures can be generated using these equations and following the code available at https://github.com/nrodgers1/Clade-dynamics-support-an-early-origin-of-crown-eukaryotes which will be deposited in a permanent repository after acceptance.

## Use of AI

Claude Ai (Anthropic) was used for grammar checks and rewording. Kimi (Moonshot AI) and Gemini (Google) were used for mathematics and code validations. No AI was used to generateoriginal data, original text, or scientific results.

## ACKNOWLEDGMENTS

This work was supported by The Leverhulme Trust grant ECF-2023-202 (C.C.F.-L.) and Human Frontier Science Program grant RGP006/2024 (N.R.). We thank two anonymous knowledgeable reviewers of an early version of this manuscript for prompting a series of revisions that significantly improved the manuscript.

## A Numerical Methods

All plotting and computation in this work is done with the standard packages NumPy^60^, SciPy^61^ and Matplotlib^62^. All plots use full versions of equations without approximations (aside from a series expansion for numerical stability when solving equation 16 for *rT* → 0). When numerically evaluating equations, we work with and plot the logarithms where possible for numerical stability. When numerically solving equation 16 we use the SciPy implementation^61^ of Brent’s Method to find the unique root. For visualisation purposes, we mask very small probabilities. In the case of figure 3, we mask probabilities below 10^*−*12^ and in figure 2 we implement masking at 10^*−*6^, which is not required in the range we plot in practice.

## B Demonstration of required properties of mean-implied gap equation

In order to demonstrate that 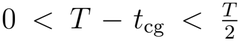 for all *r*_implied_ *>* 0 when solving equation 16, we demonstrate the left-hand side is strictly decreasing and analyse the large and small *r*_implied_ boundary limits. Firstly we write

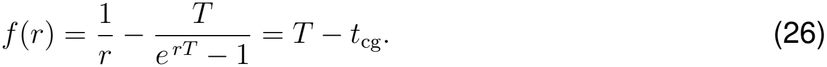

Then we make the substitution *x* = *rT*, so we have

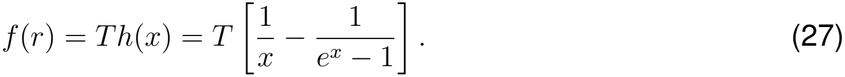

For *x* → ∞, at fixed *T*, we clearly have the gap tending to zero as the positive denominators tend to infinity. This makes sense as extremely fast diversification means the crown forms instantly. For the case *x* → 0^+^, at fixed *T*, we need to make a series expansion. We have

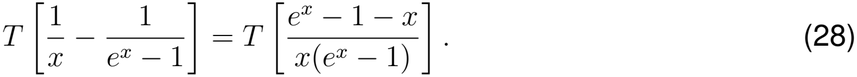

This can be expanded to second order and simplified to give

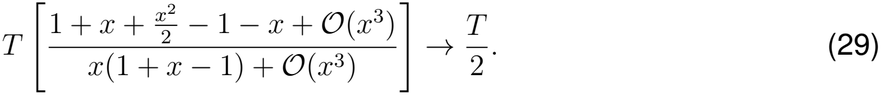

Finally, in order to show that 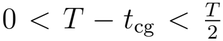 for all *r*_implied_ *>* 0, we need to show that *h*(*x*) is strictly decreasing for *x >* 0 and hence all possible gaps lie between the given extreme values. We compute

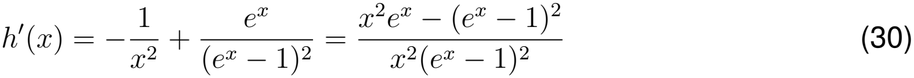

which using the positivity of the denominator implies that

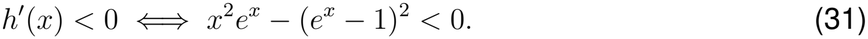

Then by expanding and multiplying by *e*^*−x*^ we have that

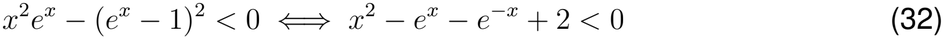

Then we use that *e*^*x*^ + *e*^*−x*^ = 2 cosh *x*, which we expand to give

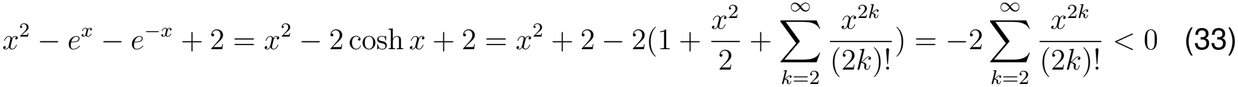

which implies that *h*^*′*^(*x*) *<* 0. Hence, this demonstrates that *h*(*x*) is strictly decreasing for *x >* 0, which in combination with the limiting values proves that 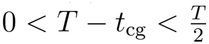 for all *r*_implied_ *>* 0. This provides a hard limit on the mean predicted length of the stem in relation to the age of a clade for diversifying clades.

